# Control of fall armyworm with Spodoptera frugiperda multiple nucleopolyhedrovirus in sweet corn cultivated under agroecological conditions

**DOI:** 10.1101/2025.07.26.666974

**Authors:** T. Cunningham, C. de la Colina, M. Pérez, G. Quintana, Marco D’Amico, M. Berretta, J Niz, M.L. Ferrelli, R. Salvador

## Abstract

The fall armyworm (FAW), *Spodoptera frugiperda* (JE Smith) is a polyphagous pest documented on more than 80 plant species. In vegetable production systems, sweet corn is their preferred host, but also infests other plants such as chilli, tomato, pumpkins, cucumber, beans and eggplant. Control strategies are commonly carried out by the use of chemical insecticides even though resistance of *S. frugiperda* populations to several insecticides has been reported. Last decades, have registered rapid adoption of organic and agroecological practices in horticulture, particularly those that rely on bioinputs such as biopesticides and biofertilizers. The baculovirus *Spodoptera frugiperda Multiple Nucleopolyhedrovirus* (SfMNPV) is a pathogenic agent for the fall armyworm and an alternative tool used for its control in sustainable pest management strategies. In this work, we evaluate two indigenous SfMNPV-based liquid formulations sprayed on sweet corn cultivated under agroecological conditions. Larvae fed with leaves treated with SfMNPV OBs showed mortality rates ranging from 55 to 68 %. Offspring from adult surviving to OBs treatments was analyzed by PCR and the results confirmed sublethal infections and vertical transmission in the fall armyworm. Corn ears harvested from each treatment were individually observed for insect presence, where the lowest rate of fall armyworm per corn ear was registered in plants treated with SfMNPV OBs. Analysis of secondary plagues in corn ears identified the sap beetle *Carpophilus dimidiatus* and the maize-infesting fly larvae *Euxesta mazorca* as the most abundant insect pests. These results show that indigenous isolates of SfMNPV produce a significant control on FAW under agroecological conditions, suggesting that the use of formulations including this virus might be a valuable tool for pest management in sustainable vegetable production systems.

**Graphical abstract:** 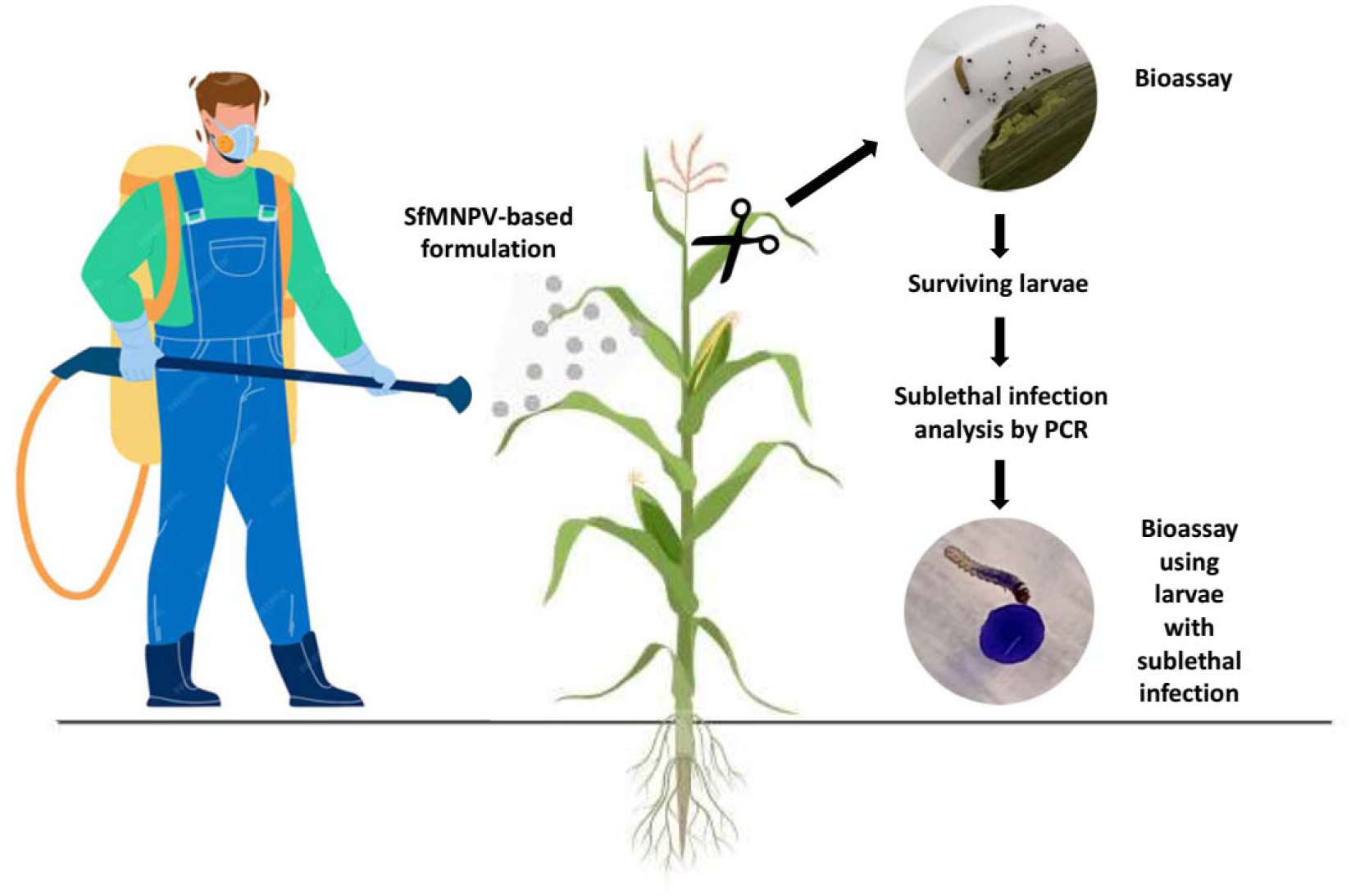

## 1. Introduction

The fall armyworm (FAW), *Spodoptera frugiperda* Smith (Lepidoptera: Noctuidae), is a native pest from tropical and subtropical regions of the Americas. In recent years, its distribution has expanded to Africa, Asia, Oceania, and most recently, Europe (Acharya et al. 2021; Babendreier et al. 2022; Tepa-Yotto et al. 2022; Wang et al. 2023). FAW larvae are polyphagous, documented on 80 plant species, including maize, rice, cotton, and pasture grasses (Montezano et al. 2018). Furthermore, it has been reported in horticultural crops such as onions, bell peppers, potatoes, tomatoes, and is particularly damaging to sweet corn (Wang et al. 2020; Acharya et al. 2022; Sisay et al. 2023). Different strategies are employed to reduce the impact of FAW on crops. The most common method involves the application of synthetic insecticides which often requires up to three treatments per season in conventional agricultural systems (Sosa et al. 2002). Due to the negative effects of chemical pesticides on the environment, their application is avoided when crops are used for human consumption. Nowadays, alternative practices are increasingly adopted in agricultural production to ensure pesticide-free crops (Finger 2024). In the last decades, agroecological methods began to be used as sustainable production that involve the elimination of chemical pesticides application (Ewert et al. 2023). A fundamental principle of agroecology emphasizes the critical roles that all organisms play in ecosystems (Wezel et al. 2020). Therefore, crop management strategies are not aimed at completely eradicating insect pests but rather at maintaining their populations density below the economic injury level. In agroecological production systems, insect pest populations can be regulated using biological control strategies (Reddy 2017). Both macro-organisms (parasitoids, predators and nematodes) and microorganisms (fungi, bacteria, viruses) are used for control pests. Members of the Baculoviridae family are a group of insect-specific viruses widely utilized as biopesticides (Hasse et al. 2015; Gelaye and Negash 2023). Baculoviruses offer numerous advantages as a tool in insect control due to their high host-specificity, typically limited to a single or a few closely related insect species, and their lack of effect on non-target organisms, including beneficial insects, plants and vertebrates (Gelaye and Negash, 2023). For decades, baculovirus-based bioinsecticides have been used in integrated pest management strategies for controlling harmful insects in agriculture (Cory and Bishop, 1997). The expanded area of organic and agroecological farming systems around the world has increased the demand of baculovirus-like biopesticides (Fenibo et al. 2022).

Several isolates of the baculovirus Spodoptera frugiperda Multiple Nucleopolyhedrovirus (SfMNPV) have been reported infecting *S. frugiperda* larvae in the Americas (Hasse et al. 2015). Additional, SfMNPV isolates were described in areas of Africa and Asia newly invaded by FAW (Hussain et al. 2021). In Argentina, the isolates SfMNPV-M and SFMNPV-C were obtained from single dead larvae collected in a maize field of the central region of the country, and in a northeastern cotton field, respectively (Berretta et al. 1998; Niz et al. 2020). Both isolates were tested in bioassays under laboratory conditions (Niz et al, 2020) and the DNA genome of SfMNPV-C was completely sequenced and the genetic content was analyzed (Masson et al., 2021). However, field trials have not yet been conducted with any of these isolates.

The field application of baculovirus insecticides requires appropriate formulations of the viral occlusion bodies (OBs), the active ingredient, along with different additives that ultimately determine their efficacy, shelf life, and ease of use (Grzywacz and Moore, 2017). The most common additives used in baculovirus formulations include surfactants, adherents, thickeners, and UV protectants (Haase et al. 2015). Several classes of baculovirus-based formulations were evaluated under field conditions. These include dry formulations, which have been applied as powders, granules, or wettable powders (Caballero et al. 2001). However, the most common method of applying baculoviral formulations is as concentrated suspensions of OBs (Haase et al. 2015). Liquid formulations are advantageous because they can be easily utilized with conventional spraying equipment. In field trials both dry and liquid SfMNPV-based were tested resulting in variable mortality rates. In Mexico and Honduras, a concentrated suspension of OBs was sprayed on plants, resulting in a low mortality rate (less than 50%), which was attributed to the insufficient persistence of OBs in the environment (Williams et al.1999). In subsequent trials, better results were obtained with improved formulations through the incorporation of UV protectants, adherents, phagostimulants and microencapsulated OBs (Belhe and Popham 2012; Gomez et al. 2013; Barrera et al. 2015).

The use of baculovirus-based formulation under field conditions may lead to the survival of some target insects due to the ingestion of a low OBs dose (Virto et al. 2017). In surviving hosts, baculovirus can be transmitted vertically resulting in sublethal infections (Ebert 2013). They are also known as covert, silent or occult infections, which are characterized by absence of visible symptoms of disease (Williams et al. 2017). Sublethal infections can be activated by different stress factors such as nutritional deficiencies, environmental changes, infections by other pathogens, parasitism or exposure to chemical agents. The activation produces a lethal viral infection, which causes the death of the insect (Williams et al. 2017). This type of baculovirus infection was described in natural populations of lepidopteran insects (Kemp and Cory 2011; Virto et al. 2014; Larem et al. 2019), and the ecological implications have not been extensively studied. Sublethal SfMNPV infections were described in *S. frugiperda* under laboratory conditions (Onkarappa et al. 2023), although the impact they could have on the management of natural populations is unknown. In this work, we evaluated two indigenous SfMNPV isolates in sweet corn cultivated at a greenhouse under agroecological conditions. In addition, we analyzed the effect of a second virus administration to FAW larvae with sublethal infection and their potential implications for pest management strategies.

## 2. Materials and Methods

### 2.1. Sources of insects and viruses

The experiments were conducted with *S. frugiperda* larvae from established colonies of insects collected in a main corn area (Santa Fe Province), and maintained at the IMYZA (INTA-Castelar). The insects were reared on an artificial diet at 25 ± 1 °C, 14:10 h light–darkness cycle, and 60% relative humidity (Greene et al. 1976). The indigenous SfMNPV isolates (SfMNPV-M and SfMNPV-C) were obtained from single dead larvae collected in a maize and cotton field of the central region of Argentina. The viruses were propagated on their original hosts and OBs were purified as previously described (Berretta et al. 1998). The OBs doses used in treatments were estimated based on lethal dose 50 (Niz et al. 2015).

### 2.2 Evaluation of liquid SfMNPV OBs formulation

A liquid formulation with a minimum of components was designed and evaluated. A suspension containing SfMNPV OBs as the active ingredient, the mixture also included a commercial polyethylene glycol-based surfactant/adhesive and a biologically derived carminic acid as UV protectant. To determine whether the mix of these compounds were inherently toxic to S. frugiperda larvae or could alter the insecticidal capacity of SfMNPV OBs, tests were conducted with the formulation on third and four instar larvae. Thus, three treatments were evaluated: a mixture of baculovirus SfMNPV-M (7.3 × 10^5^ OBs/ml), surfactant/adhesive, UV protectant, 1% sucrose, and 0.1% Coomasie Blue (Treatment 1); a mixture containing surfactant/adhesive, UV protectant, 1% sucrose, and 0.1% Coomasie Blue (Treatment 2); and a mixture containing 1% sucrose and 0.1% Coomassie Blue (Treatment 3). Three replicates of 25 larvae each were prepared for each treatment. The solutions were administered using the droplet feeding technique (Hughes and Wood, 1981). Mortality was recorded at 24-hour intervals. Dead larvae were collected and kept at −20°C. Statistical analysis was performed using the methodology described by Niz et al. (2020).

### 2.3 Greenhouse trial

A liquid SfMNPV formulation was evaluated on sweet corn plants cultivated in the greenhouse at Gorina Experimental Station (34° 54’ 51.890” S, 58° 2’ 17.683” W). Sweet corn seeds (variety Anita-INTA) were planted individually at 40 cm from each other in five rows of 25 m and 1,20 m between rows. Likewise, each row was subdivided in five sections corresponding to five treatments. Each section included a total of 20 plants. A greenhouse trial was performed in February with median temperatures ranging between 25-28 °C. Plants of 1,80 m with two ears were sprayed between 7:00 and 8:00 am local time. The treatments were applied using a backpack sprayer and the volume sprayed to each plant was on average 20 ml, most of which was directed to leaves and corn ears. A total of five treatments with three replicates were evaluated: a negative control group without treatment (Treatment 1), a positive control was applied containing the bioinsecticide “Bacthur” (Ando and Co.) based on entomopathogenic bacteria *Bacillus thuringiensis* (Treatment 2), a formulation including surfactant/adhesive and UV protector but without OBs (Treatment 3), a formulation including surfactant/adhesive, UV protector, and 7,3×10^5^ OBs/mL of SfMNPV-M isolate (Treatment 4), and a formulation including surfactant/adhesive, UV protector, and 5×10^5^ OBs/mL of SfMNPV-C isolate (Treatment 5). At 2 hours after treatment application a total of 10 recently emerged neonate larvae were distributed for each plant.

The evaluation of SfMNPV virus-based formulations was performed using the methodology described by Behle et al. (2012). Leaf samples of each treatment were collected two hours after spraying. A total of thirty leaf discs (2 cm of diameter) were cut from each treatment (with three replicates) and used individually to feed third instar FAW larvae during 24 hs, then each larva was transferred to semisynthetic diet (Green et al.1976) and reared in laboratory under controlled conditions. Mortality was recorded at 24-hour intervals for two weeks. Dead larvae were collected individually and preserved at −20°C. Mortality data were statistically analyzed according to Niz et al. (2020). An additional evaluation was performed documenting the presence of fall armyworm larvae on each corn ear in all plants. Thus, total ears from each treatment were harvested at two weeks post spraying and the presence of larvae was individually analyzed and recorded. Moreover, the presence of other insects such as beetles, dipteran larvae, and caterpillars was documented.

### 2.4. PCR analysis

Dead larvae registered during laboratory assay were individually preserved at −20°C and processed to determine the presence of SfMNPV DNA using the Polymerase Chain Reaction (PCR) technique. DNA samples were prepared as described by Manzan et al. (2008). The presence of SfMNPV DNA in treated and control larvae was determined using specific primers designed to amplify the SfMNPV ORF 44 in both isolates (Masson et al. 2021). PCRs were performed using a thermal cycler model T18 (Ivema Brand). The reaction mixture consisted of: 1.5 mM MgCl2 (0.3 µl); 2.5 mM dNTPs (1 µl); 0.5 U of DNA Taq polymerase (0.3 µl) with an optimal elongation temperature of 72°C (Inbio Highway) and 0.5 µM of each primer (0.6 µl), in a final volume of 10 µl. The cycling was as follows: 1. Initial denaturation (94°C, 4 min); 2. Denaturation (94°C, 2 min); 3. Annealing (53°C, 40 sec), 4. Extension (72°C, 40 sec), steps 2 to 4 were repeated 39 times, followed by a final extension (72°C, 3 min). The PCR products were analyzed through horizontal electrophoresis using 1% agarose gels in TAE buffer (Tris-HCl, acetic acid, EDTA). GelRed (Biotum Co.) was used as agent for visualization.

### 2.5. Sublethal infections analysis

The surviving larvae from field evaluation of SfMNPV virus-based formulations and controls were maintained with artificial diet to adult stage. Larval progeny in the second stage was individually analyzed by PCR to detect SfMNPV. Total DNA was individually extracted from larvae (whole body) following a CTAB protocol and stored at −20∘C until PCR analysis. PCRs were performed following the methodology described above. To evaluate the effect of administration of SfMNPV on FAW larvae with sublethal infection a bioassay was carried out by feeding OBs to newly molted 3rd instar larvae, after 12 hr of starvation, using the droplet-feeding method described by Hughes and Wood (1981). The experiments were replicated three times and larval mortality was recorded for 2 weeks. Mortality data were statistically analyzed according by Niz et al. (2020).

## 3. Results

### 3.1. Evaluation of SfMNPV-liquid formulation under laboratory conditions

*Spodoptera frugiperda* larvae were fed with liquid suspension containing SfMNPV OBs, a surfactant and a UV protectant (Treatment 1). Control groups with solution with surfactant and a UV protectant (Treatment 2), and untreated larvae (Treatment 3) were evaluated. Third instar larvae fed with OBs suspension showed a statistically significant mortality rate (80 %) compared with control treatments (Figure 1, left). Identical OBs suspension produced a lower mortality on the fourth stage of FAW (35 %) but differences with control treatments were still statistically significant (Figure 1, right). SfMNPV mortalities were similar to previous bioassays performed by our group and the addition of surfactant/adhesive and UV protectant did not contribute to mortality.

**Figure 1.**
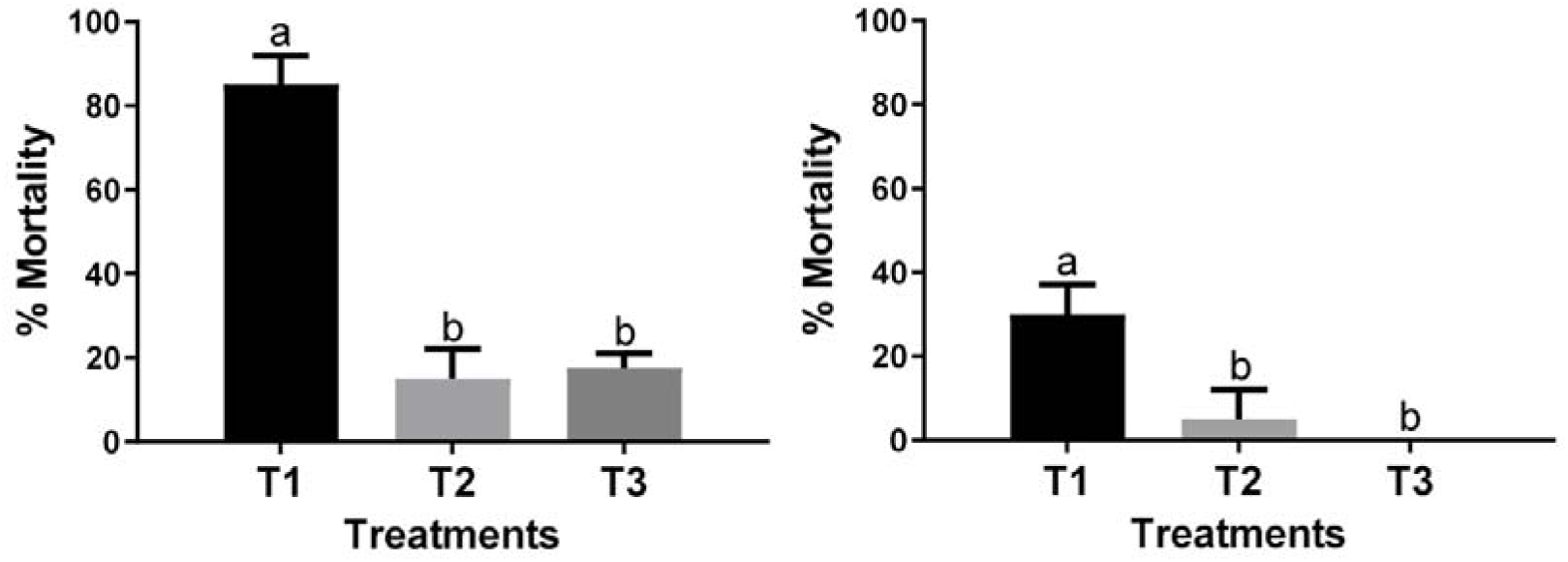
Mortality registered in *S. frugiperda* larvae fed with formulated SfMNPV OBs. **Left:** 3rd instar *S. frugiperda*. **Right:** 4th instar *S. frugiperda.* T1: SfMNPV OBs, surfactant and UV protectant. T2: surfactant and UV protectant. T3: larvae untreated larvae. Same letter was not significantly different (p<0.05).

### 3.2. Test of SfMNPV-based formulation at greenhouse

We then tested the formulation with both isolates M and C under greenhouse conditions. Third instar larvae of *Spodoptera frugiperda* were fed with sprayed leaf corn disks and the final mortalities were registered (Figure 2, left). The highest mortality rate was observed in exposed larvae to formulated SfMNPV-C (68%, T5). The treatment with SfMNPV-M showed lower mortality (55%, T4) compared with SfMNPV-C, but the difference between both was not significantly different. Also, the mortality rates of SfMNPV treatments were not significantly different to those registered in *Bt* treatment (47%, T3). However, the scored mortalities in both SfMNPV and *Bt* treatments were significantly higher than negative control treatments (T1, T2).

**Figure 2.**
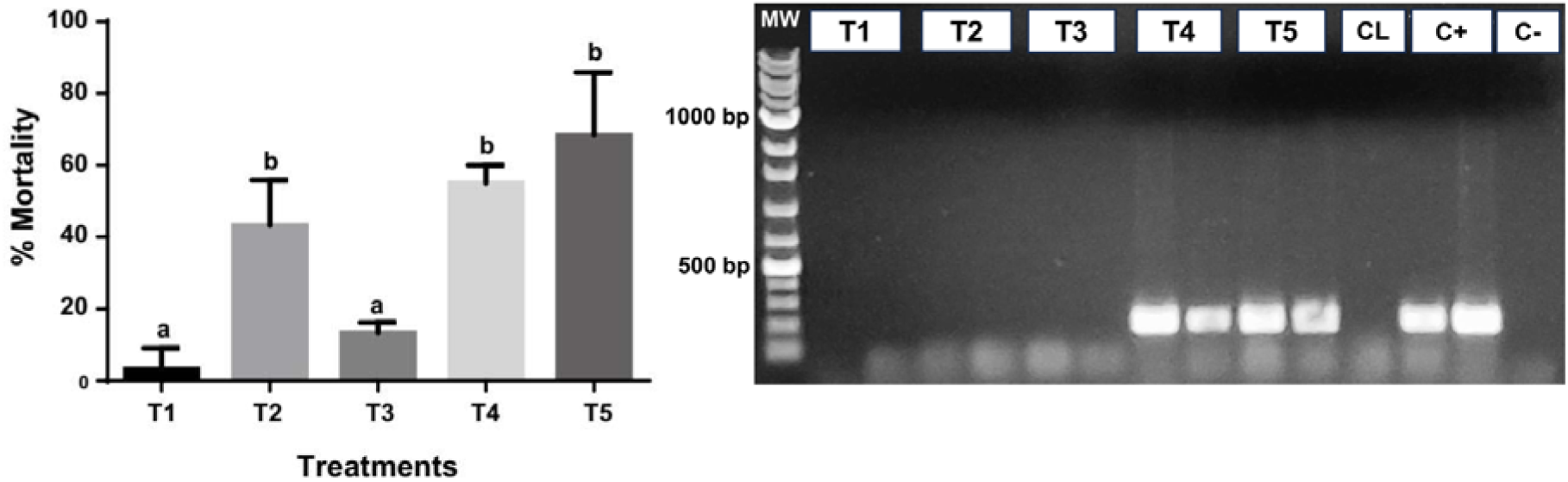
**Left: Mortality registered in *S. frugiperda* larvae under different treatments**. T1: Negative control group without treatment. T2: Positive control with *Bacillus thuringiensis*. T3: Formulation control without virus. T4: Formulation with SfMNPV-M. T5: Formulation with SfMNPV-C. Treatments with the same letter were not significantly different (p<0.05). **Right: Detection of SfMNPV DNA in *S. frugiperda* larvae**. MW: molecular weight (1 kb. Inbio Highway Co T1: Negative control group without treatment. T2: Positive control with *Bacillus thuringiensis*. T3: Formulation control without virus. T4: Formulation with SfMNPV-M. T5: Formulation with SfMNPV-C. CL: negative control (uninfected larva); C+: positive control I (SfMNPV DNA) positive control II (plasmid with fragment of SfMNPV 44 ORF); C-: PCR negative control

In order to confirm that dead larvae registered in bioassay were due to the SfMNPV baculovirus, total genomic DNA was extracted from whole larval body and analyzed it by PCR using specific SfMNPV primers corresponding to ORF 44 (Figure 2, right). The presence of viral DNA only was detected in larvae fed with foliar disc from treatments T4 and T5.

### 3.3. Presence of *S. frugiperda* larvae in corn ears

Under high densities of fall armyworm infestations in maize crops, larvae can burrow into the terminal end of the ear (the silk end) leading to increased grain loss. To assess ear damage caused by *S. frugiperda* larvae, all corn ears were harvested from each treatment and analyzed to detect the presence of larvae (Figure 3). The results showed significant differences between plants treated with SfMNPV based-formulation (T4 and T5) compared to other treatments. Plants treated with Bt (T2) did not exhibit significant differences with untreated groups (T1) and plants treated with the formulation without virus (T3). The presence of fall armyworm larvae in treatments T1, T2, and T3 ranged between 30% and 40% of analyzed corn ears, while for the treatments with baculovirus, T4 and T5, the proportion of ears with insect presence was considerably lower, with larvae found in 15% of ears.

**Figure 3.**
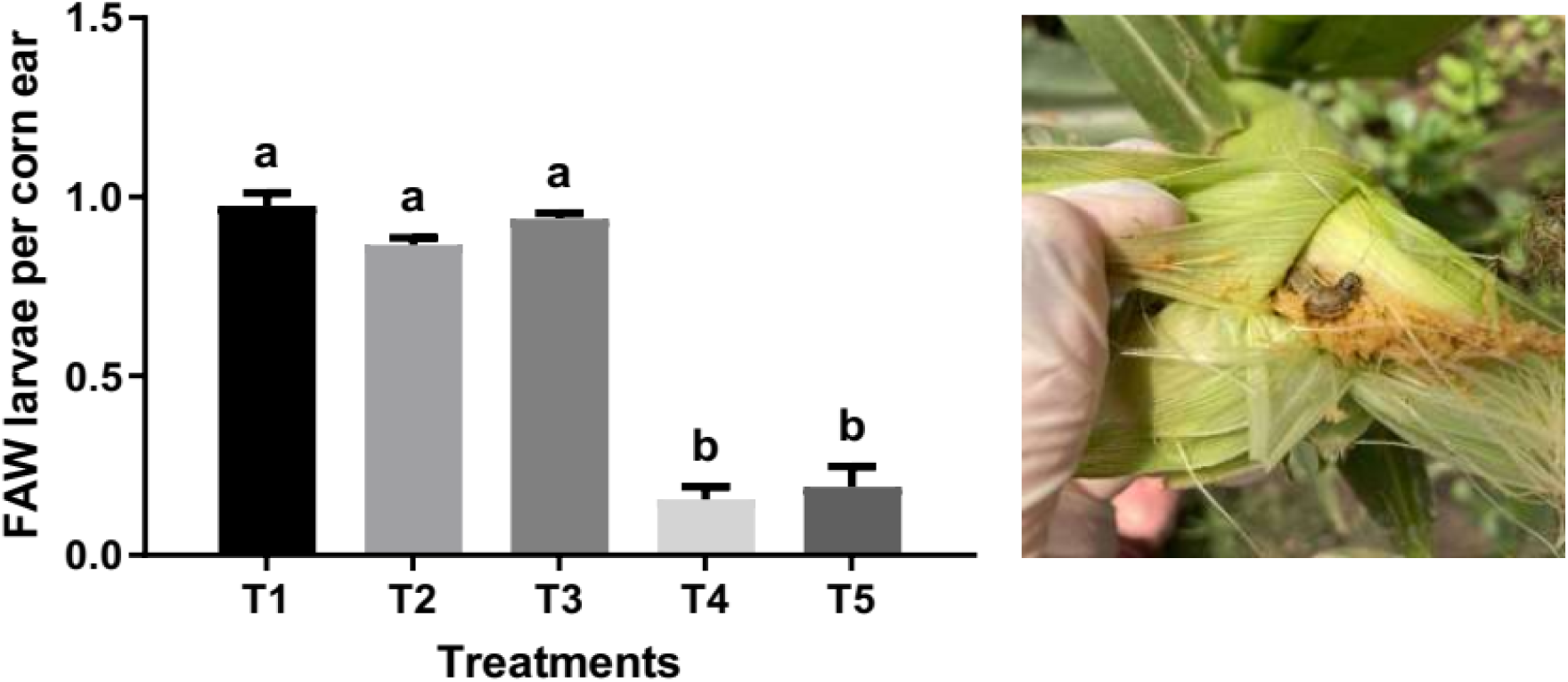
Larvae registered per corn ear. T1: plants without treatment. T2: treatment with Bacillus thuringiensis. T3: control of formulated without viruses. T4: formulated with SfMNPV-M. T5: formulated with SfMNPV-C. Treatments with the same letter was not significantly different (p<0.05)

### 3.4. Presence of others pests in corn ears

The abundance of insects on corn ears was assessed both by treatment and across the entire trial. Statistical analysis of treatment effects revealed no significant differences in insect presence among the experimental groups (Supplementary material). Regarding species-specific prevalence in the entire trial (Figure 4. Left), the sap beetle *Carpophilus dimidiatus* (Fabricius) exhibited the highest incidence, affecting 46% of sampled corn ears. The larval stage of the corn-infesting fly *Euxesta* mazorca was detected on 44% of ears, while ants *Acromyrmex lundi* and the lepidopteran *Helicoverpa* sp. occurred at lower frequencies (8% and 1%, respectively). A small proportion of sampled corn ears exhibited co-infestation by FAW and another insect (Figure 4. Right). The most frequent interspecific association involved FAW with fly larvae (*Euxesta mazorca*) and sap beetles (*Carpophilus dimidiatus*), occurring in 25% of analyzed corn ears.

**Figure 4.**
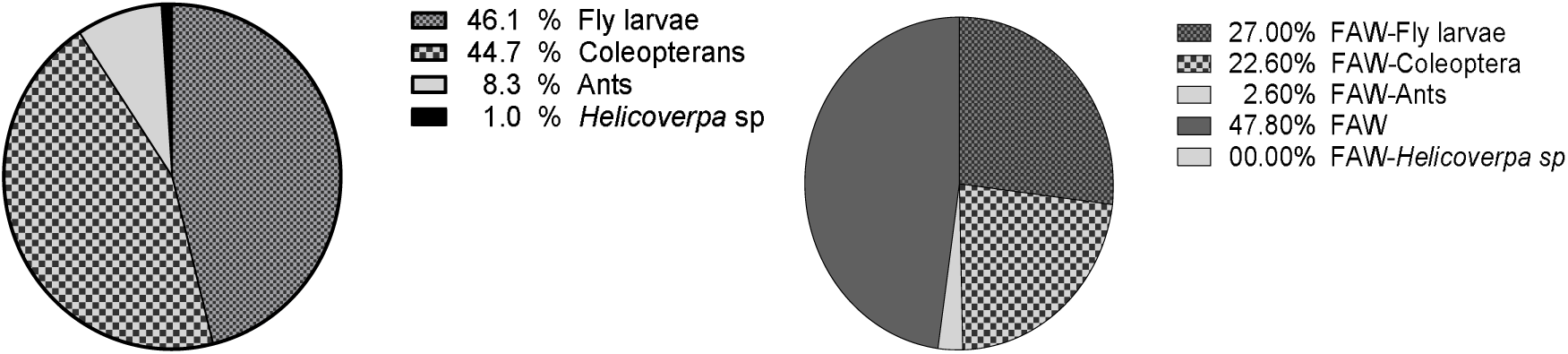
Detection of insects in total corn ears. **Left:** Insects presence in total corn ears analyzed. **Right**: Corn ear with co-infestation by FAW and another insect.

### 3.5. Analysis of sublethal SfMNPV infection on FAW

In order to analyze the persistence of initial SfMNPV input we tested surviving larvae for virus presence. Surviving larvae from treatments T4 and T5 of the bioassay described above were maintained in laboratory conditions to the adult stage. Their larval progeny was analyzed by PCR to detect inherited sublethal SfMNPV infection. The results demonstrated that the virus was detected in all insects analyzed (Figure 5, left). Then, we infected these larvae per os to evaluate if new OB input triggered a lethal infection. Third-instar larvae with confirmed sublethal infection were treated with SfMNPV OBs, and mortality was recorded daily (Figure 5, right). The mortality rate recorded in larvae with sublethal infection fed with SfMNPV OBs was 80 %, a similar value to that registered in bioassay performed with healthy larvae (Figure 1).

**Figure 5.**
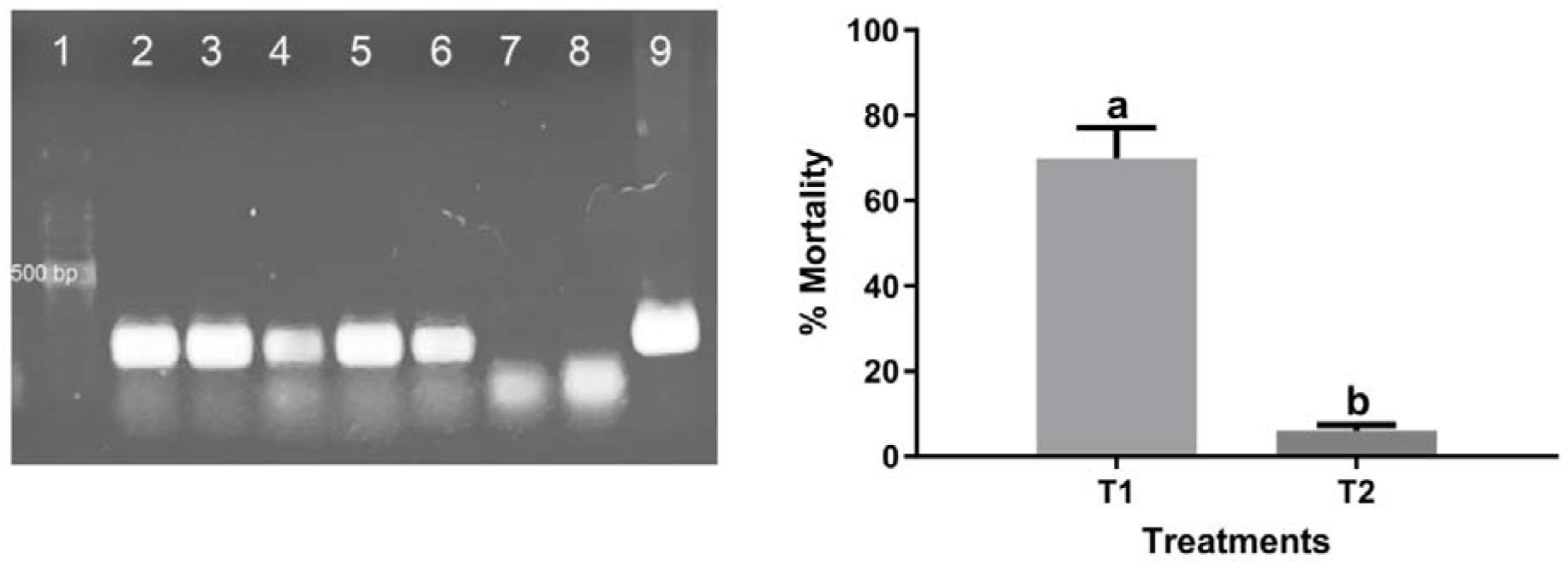
Sublethal SfMPV infection in FAW larvae. **Left. Detection of SfMNPV DNA by PCR**. Line 1: molecular weight (1 kb. Inbio Highway Co); Lines 2-6: individual larva; Lines 7: control larva; Lines 8: negative control: Lines 9: positive control (SfMNPV DNA). **Right. Bioassay with sublethal-infected FAW larvae.** T1: FAW larvae with sublethal infection fed with SfMNPV OBs solution. T2: FAW larvae with sublethal infection fed with control solution (Sugar and colorant). Different letters indicate significant differences (Annex 4). Treatments with the same letter was not significantly different (p<0.05)

## 4. Discussion

In recent decades, fresh food production systems have transitioned toward sustainable agricultural practices with reduced dependence on synthetic agrochemicals (Zhang 2024). Within this paradigm shift, baculovirus-based biopesticides have emerged as increasingly prominent tools for effective pest management, demonstrating particular efficacy against lepidopteran crop pests (Hasse et al. 2015; Martinez-Belardi et al. 2025). In our study, we evaluated SfMNPV-based baculovirus formulations in an agroecological production system, achieving target pest control efficacy ranging from 55% to 68% depending on the viral isolate employed. These findings are consistent with previous studies investigating SfMNPV formulations under different methodological approaches and production systems (Williams et al. 1998; Martinez et al. 2000; Behle et al. 2012; Cuartas-Otalora et al. 2019). In our greenhouse trial, no chemical control methods were applied against other pest species observed on the plants. Post-harvested analysis of corn ears revealed the presence of other common pests in the crop such as the beetle (*Carpophilus dimidiatus)*, fly larvae (*Eusexta mazorca)*, ants (*Acromyrmex lundi)* and corn earworm (*Helicover* zea). The incidence of these pests in maize appeared to be independent of the treatments applied in the present work. Similar insect species were reported by Williams et al. (1998) in their study, although unlike our trials, they recorded an increase in these insects in SfMNPV OBs treatments; this could be due to the use of sugar-containing formulations which could have favored the attraction or survival of non-target insects. It is also important to mention that observed low incidence of *Helicoverpa* sp. larvae, typically common in corn ears, may be attributed to interspecific competition with fall armyworm. Song et al. (2024) demonstrated in a recent work that FAW’s ecological dominance over *Helicoverpa* spp. can be explained by both its aggressive larval behavior and predatory advantages during larval stages.

Many studies indicated that baculoviruses can establish sublethal or covert infections in insects, characterized by viral persistence without causing the host mortality (Burden et al. 2002; Matthews et al. 2002; Cabodevila et al. 2011; Larem et al. 2019). In our study, a sublethal infection was detected in live FAW larvae previously treated with SfMNPV. PCR analysis on progeny from these larvae confirmed virus presence, indicating a vertical transmission. These results corroborate previous findings reported for an Indian isolate of SfMNPV, which demonstrated the ability to establish covert infections in *Spodoptera frugiperda*, with evidence of vertical transmission to offspring (Onkarappa et al. 2023). Sublethal baculovirus infections produce on the host insect a significant reduction in pupal weight, adult fecundity and fertility (Myers et al. 2002, Vilaplana et al. 2010). Furthermore, previous studies performed on *Spodoptera exigua* have demonstrated that sublethally infected insects may exhibit an increased susceptibility to reinfection by occlusion bodies of the same viral isolate compared to uninfected insects (Cabodevila et al. 2011). Thus, from a pest control point of view, an insect population with sublethal infection generated after field application could be beneficial for pest management in subsequent generations (Williams et al. 2017). In our bioassays, results showed similar mortality rates between primarily infected FAW larvae and those with established sublethal SfMNPV infections. These findings are consistent with those of Virto et al. (2017), who reported that progeny from greenhouse-insects treated with OBs showed no significant difference in susceptibility to reinfection by the homologous viral isolate. Further experiments are required to determine final susceptibility of SfMNPV reinfection on *Spodoptera frugiperda* larvae.

From an agroecological perspective, insects are considered essential components of the agricultural systems (Jeanneret et al. 2021; Steffan-Dewenter et al. 2024). Although herbivorous insects are frequently regarded as agricultural pests, they fulfill an essential ecological function in agroecosystems by serving as a food resource for natural enemies including predatory arthropods, parasitoids, and insectivorous vertebrates such as birds and small mammals. Consequently, agroecological management strategies prioritize regulating pest insect population densities rather than complete eradication. The global shift toward pesticide-free fresh food production systems demands the development and deployment of effective, ecologically sustainable biological control agents (Jacquet et al. 2022). The results presented in this work demonstrate that the two indigenous SfMNPV-based bioinsecticides are capable of achieving significant control of FAW and could establish itself in the insect population through sublethal infections under agroecological conditions.

## 5. Declarations

### Ethical Approval

This article does not contain any studies with human participants or animals performed by any of the authors.

### Conflicts of Interest

The authors declare no conflict of interest.

### Funding

This research was funded by grants from Instituto Nacional de Tecnología Agropecuaria (INTA): Proyect I073 and Ministerio de Producción, Ciencia e Innovación Tecnológica de Buenos Aires: FITBA 2022-B16 to R. Salvador.

## Acknowledgments

We thank Marcelo Farinon (IMyZA-INTA) and Lorena Ruiz Diaz (IMyZA-INTA) for the maintenance of the insect colony at INTA. Vanesa Castaldo, Jorge Luna and Gorina Experimental Station Staff are gratefully acknowledged for their assistance in greenhouse trial.

## Authors’ contributions

RS, TC and CDC conceived and designed the laboratory experiments. JN, MFB and RS participated in viral amplification and purification from insect larvae. RS, TC and GQ participated in SfMNPV formulation. RS, MP y MD conceived and designed greenhouse trial. RS, LF, TC, CDC, MP and MD collaborated in greenhouse trial. RS wrote the main manuscript text. All authors review and approved the final manuscript.

## Supplementary material

**Supplementary material 1.**
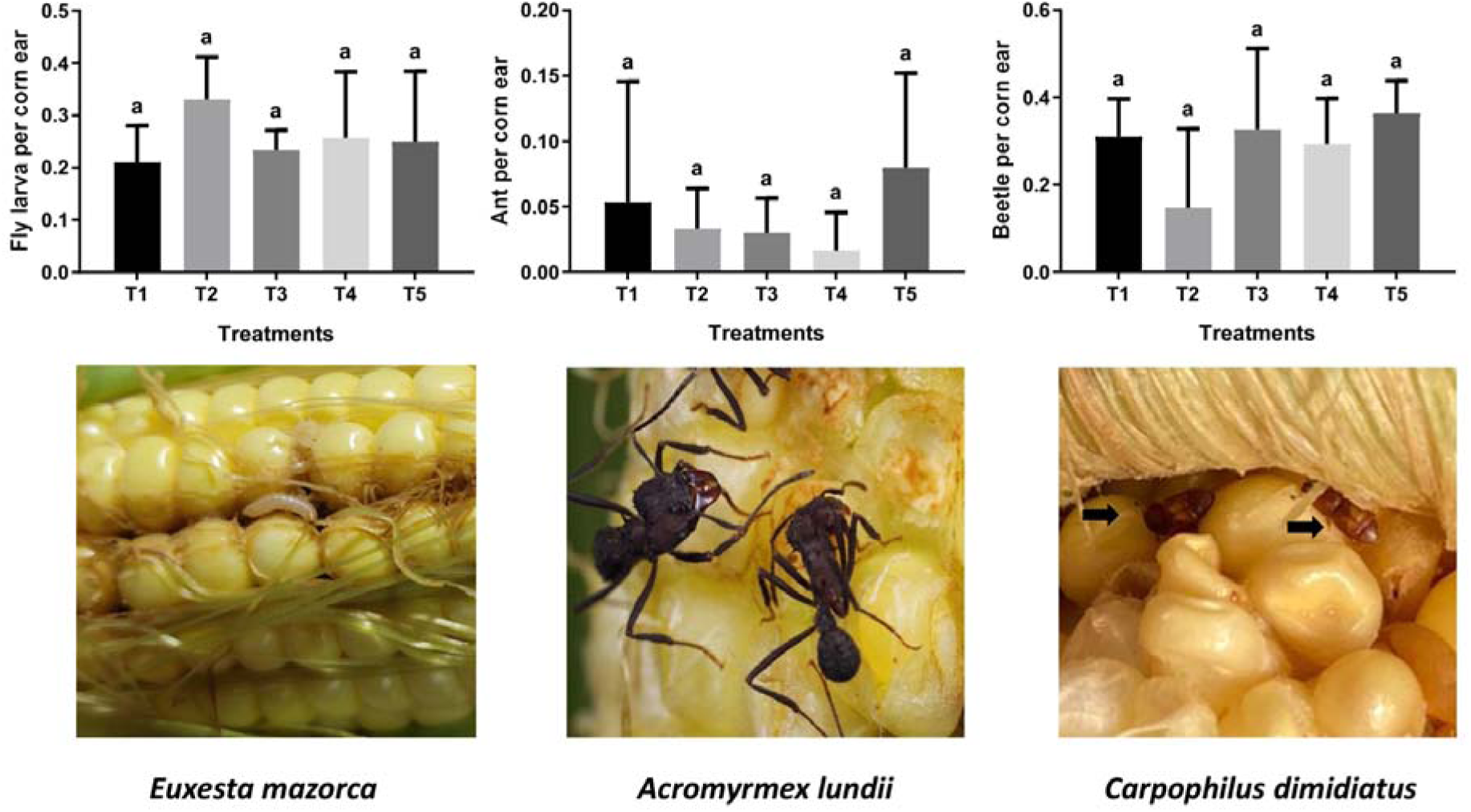
Insect presence by treatments. T1: plants without treatment. T2: treatment with Bacillus thuringiensis. T3: control of formulated without viruses. T4: formulated with SfMNPV-M. T5: formulated with SfMNPV-C. Treatments with the same letter was not significantly different (p<0.05).

## Notes

### Competing Interest Statement

The authors have declared no competing interest.

## References

Acharya R, et al. (2021). Genetic Relationship of Fall Armyworm (Spodoptera frugiperda) Populations That Invaded Africa and Asia. Insects. 2021 May 12;12(5):439. doi: 10.3390/insects12050439. PMID: 34066149; PMCID: PMC8151712.

Behle RW, Popham HJ. (2012). Laboratory and field evaluations of the efficacy of a fast-killing baculovirus isolate from Spodoptera frugiperda. J Invertebr Pathol. Feb;109(2):194–200. doi: 10.1016/j.jip.2011.11.002. Epub 2011 Nov 12. PMID: 22100417.

Burden JP, Griffiths CM, Cory JS, Smith P, Sait SM. (2002). Vertical transmission of sublethal granulovirus infection in the Indian meal moth, Plodia interpunctella. Mol Ecol. Mar;11(3):547–55. doi: 10.1046/j.0962-1083.2001.01439.x. PMID: 11918789.

Caballero, P., López-Ferber, M. & Williams, T. (2001) Los baculovirus y sus aplicaciones como bioinsecticidas en el control biológico de plagas. pp. 518. Phytoma S.A., Valencia, España (ISBN: 84-932056-0-5).

Cabodevilla O, Villar E, Virto C, Murillo R, Williams T, Caballero P. (2011). Intra- and ntergenerational persistence of an insect nucleopolyhedrovirus: adverse effects of sublethal disease on host development, reproduction, and susceptibility to superinfection. Appl Environ Microbiol. May;77(9):2954–60. doi: 10.1128/AEM.02762-10. Epub 2011 Mar 11. PMID: 21398487; PMCID: PMC3126408.

Cory JS, Bishop DH. (1997). Use of baculoviruses as biological insecticides. Mol Biotechnol. 1997 Jun;7(3):303–13. doi: 10.1007/BF02740821. PMID: 9219244.

Ewert, F., Baatz, R., & Finger, R. (2023). Agroecology for a sustainable agriculture and food system: from local solutions to large-scale adoption. Annual Review of Resource Economics, 15(1), 351–381.

Fenibo, E.O., Ijoma, G.N., Matambo, T. (2022). Biopesticides in Sustainable Agriculture: Current Status and Future Prospects. In: Mandal, S.D., Ramkumar, G., Karthi, S., Jin, F. (eds) New and Future Development in Biopesticide Research: Biotechnological Exploration. Springer, Singapore. 10.1007/978-981-16-3989-0_1

Finger, Robert. (2024). On the definition of pesticide-free crop production systems, Agricultural Systems, Volume 214, 103844, ISSN 030821X, 10.1016/j.agsy.2023.103844.

Gelaye, Y., & Negash, B. (2023). The role of baculoviruses in controlling insect pests: A review. Cogent Food & Agriculture, 9(1). 10.1080/23311932.2023.2254139

Gómez, J.; Guevara, J.; Cuartas, P.; Espinel, C.; Villamizar, L. (2013). Microencapsulated Spodoptera frugiperda nucleopolyhedrovirus: Insecticidal activity and effect on arthropod populations in maize. Biocontrol Sci. Technol. 23, 829–847.

Greene GL, Leppla NC, Dickerson WA (1976) Velvetbean caterpillar: a rearing procedure and artificial medium. J Econ Entomol 69:487–488

Grzywacz, D and Moore, S. (2017). Chapter 7. Production, Formulation, and Bioassay of Baculoviruses for Pest Control, Editor(s): Lawrence A. Lacey, Microbial Control of Insect and Mite Pests, Academic Press, Pages 109–124, ISBN 9780128035276, 10.1016/B978-0-12-803527-6.00007-X.

Haase S, Sciocco-Cap A, Romanowski V. (2015). Baculovirus insecticides in Latin America: historical overview, current status and future perspectives. Viruses. Apr 30;7(5):2230–67. doi: 10.3390/v7052230. PMID: 25941826; PMCID: PMC4452904.

Hughes, P.R., Wood, H.A. (1981). A synchronous peroral technique for the bioassay of insect viruses. J. Invertebr. Pathol. 37, 154–159.

Hussain AG, Wennmann JT, Goergen G, Bryon A, Ros, Vid. (2021). Viruses of the Fall Armyworm Spodoptera frugiperda: A Review with Prospects for Biological Control. Viruses. Nov 4;13(11):2220. doi: 10.3390/v13112220. PMID: 34835026; PMCID: PMC8625175.

Jacquet, F., Jeuffroy, MH., Jouan, J. et al. (2022). Pesticide-free agriculture as a new paradigm for research. Agron. Sustain. Dev. 42, 8. 10.1007/s13593-021-00742-8

Jeanneret P, Aviron S, Alignier A, Lavigne C, Helfenstein J, Herzog F, Kay S, Petit S. (2021). Agroecology landscapes. Landsc Ecol. 36(8):2235–2257. doi: 10.1007/s10980-021-01248-0. Epub 2021 Jun 26. PMID: 34219965; PMCID: PMC8233588.

Kemp EM, Woodward DT, Cory JS. (2011). Detection of single and mixed covert baculovirus infections in eastern spruce budworm, Choristoneura fumiferana populations. J Invertebr Pathol. Jul;107(3):202–5. doi: 10.1016/j.jip.2011.05.015. Epub 2011 May 15. PMID: 21616077.

Larem A, Ben Tiba S, Fritsch E, Undorf-Spahn K, Wennmann JT, Jehle JA. (2019). Effects of a Covert Infection with Phthorimaea operculella granulovirus in Insect Populations of Phthorimaea operculella. Viruses. Apr 9;11(4):337. doi: 10.3390/v11040337. PMID: 30970670; PMCID: PMC6520744.

Manzan, M.A., Aljinovic, E.M., Biedma, M.E., Sciocco-Cap, A., Ghiringhelli, P.D., Romanowski, V., (2008). Multiplex PCR and quality control of Epinotia aporema granulovirus production. Virus Genes 37, 203–211.

Martínez-Balerdi, M., Caballero, J., Aguirre, E., Caballero, P., & Beperet, I. (2025). Baculoviruses as Microbial Pesticides: Potential, Challenges, and Market Overview. Viruses, 17(7), 917. 10.3390/v17070917

Masson T, Fabre ML, Pidre ML, Niz JM, Berretta MF, Romanowski V, Ferrelli ML. (2021) Genomic diversity in a population of Spodoptera frugiperda nucleopolyhedrovirus. Infect Genet Evol. Jun; 90:104749. doi: 10.1016/j.meegid.2021.104749. Epub 2021 Feb 2. PMID: 33540087.

Matthews HJ, Smith I, Edwards JP. (2002). Lethal and sublethal effects of a granulovirus on the tomato moth Lacanobia oleracea. J Invertebr Pathol. Jun;80(2):73–80. doi: 10.1016/s0022-2011(02)00100-3. PMID: 12383432.

Migliorini, P., Wezel, A. (2017). Converging and diverging principles and practices of organic agriculture regulations and agroecology. A review. Agron. Sustain. Dev. 37, 63. 10.1007/s13593-017-0472-4

Myers J. H., Malakar R., Cory J. S. (2000). Sublethal nucleopolyhedrovirus infection effects on female pupal weight, egg mass size and vertical transmission in gypsy moth (Lepidoptera: Lymantriidae). Environ. Entomol. 29:1268–1272

Nagoshi RN, Meagher RL. (2022). The Spodoptera frugiperda Host Strains: What They Are and Why They Matter for Understanding and Controlling This Global Agricultural Pest. J Econ Entomol 115:1729–1743. 10.1093/jee/toac050

Reddy, P.P. (2017). Agro-Ecological Pest Management – An Overview. In: Agro-ecological Approaches to Pest Management for Sustainable Agriculture. Springer, Singapore. 10.1007/978-981-10-4325-3_1

Sisay B, Sevgan S, Weldon CW, Krüger K, Torto B, Tamiru A. (2023). Responses of the fall armyworm (Spodoptera frugiperda) to different host plants: Implications for its management strategy. Pest Manag Sci. Feb;79(2):845–856. doi: 10.1002/ps.7255. Epub 2022 Nov 18. PMID: 36301535.

Song, Y.; Li, H.; He, L.; Zhang, H.; Zhao, S.; Yang, X.; Wu, K. (2023). Interspecific Competition between Invasive *Spodoptera frugiperda* and Indigenous *Helicoverpa armigera* in Maize Fields of China. Agronomy, 13, 911. 10.3390/agronomy13030911

Tepa-Yotto GT, Chinwada P, Rwomushana I, Goergen G, Subramanian S. (2022). Integrated management of *Spodoptera frugiperda* 6 years post detection in Africa: a review. Curr Opin Insect Sci. Aug;52:100928. doi: 10.1016/j.cois.2022.100928. Epub 2022 May 6. PMID: 35534003.

Vilaplana L., Kenneth W., Redman E. M., Cory J. S. (2010). Pathogen persistence in migratory insects: high levels of vertically-transmitted virus infection in field populations of the African armyworm. Evol. Ecol. 24:147–160

Virto C, Navarro D, Tellez MM, Herrero S, Williams T, Murillo R, Caballero P. (2014). Natural populations of Spodoptera exigua are infected by multiple viruses that are transmitted to their offspring. J Invertebr Pathol. Oct; 122:22–7. doi: 10.1016/j.jip.2014.07.007. Epub 2014 Aug 13. PMID: 25128697.

Virto, C., Williams, T., Navarro, D. et al. (2017). Can mixtures of horizontally and vertically transmitted nucleopolyhedrovirus genotypes be effective for biological control of Spodoptera exigua?. J Pest Sci 90, 331–343. 10.1007/s10340-016-0743-x

Wang J, Huang Y, Huang L, Dong Y, Huang W, Ma H, Zhang H, Zhang X, Chen X, Xu Y. (2023). Migration risk of fall armyworm (Spodoptera frugiperda) from North Africa to Southern Europe. Front Plant Sci. Apr 3;14: 1141470. doi: 10.3389/fpls.2023.1141470. PMID: 37077648; PMCID: PMC10106561.

Wang W, He P, Zhang Y, Liu T, Jing X, Zhang S. (2020). The Population Growth of Spodoptera frugiperda on Six Cash Crop Species and Implications for Its Occurrence and Damage Potential in China. Insects. Sep 17;11(9):639. doi: 10.3390/insects11090639. PMID: 32957580; PMCID: PMC7563954.

Wezel, A., et al. (2020). Agroecological principles and elements and their implications for transitioning to sustainable food systems. A review. Agron. Sustain. Dev. 40, 40. 10.1007/s13593-020-00646-z

Zhang QF. (2024). From Sustainable Agriculture to Sustainable Agrifood Systems: A Comparative Review of Alternative Models. Sustainability. 16(22):9675. 10.3390/su16229675

